# Widespread shortening of 3’ untranslated regions and increased exon inclusion are evolutionarily conserved features of innate immune responses to infection

**DOI:** 10.1101/026831

**Authors:** Athma A. Pai, Golshid Baharian, Ariane Pagé Sabourin, Jessica F. Brinkworth, Yohann Nédélec, Joseph W. Foley, Jean-Christophe Grenier, Katherine J. Siddle, Anne Dumaine, Vania Yotova, Zachary P. Johnson, Robert E. Lanford, Christopher B. Burge, Luis B. Barreiro

## Abstract

The contribution of pre-mRNA processing mechanisms to the regulation of immune responses remains poorly studied despite emerging examples of their role as regulators of immune defenses. Here, we used mRNA sequencing to quantify gene expression and isoform abundances in primary macrophages from 60 individuals, before and after infection with two live bacteria. In response to both bacteria we identified thousands of genes that significantly change isoform usage in response to infection, and found global shifts towards (*i*) the inclusion of cassette exons and (*ii*) shorter 3’ UTRs. Using complementary data collected in non-human primates, we show that these features are evolutionarily conserved among primates. Finally, our results suggest that the pervasive usage of shorter 3’ UTRs is a mechanism for particular genes to evade repression by immune-activated miRNAs. Collectively, our results show that dynamic changes in RNA processing play a key role in the regulation of innate immune responses.

## INTRODUCTION

Innate immune responses depend on robust and coordinated gene expression programs involving the transcriptional regulation of thousands of genes (Huang et al. 2001; Smale 2010; Medzhitov and Horng 2009). These regulatory cascades start with the detection of microbial-associated products by pattern recognition receptors, which include Toll Like Receptors (TLRs), NOD-like receptors and specific C-type lectins (Medzhitov and Janeway 1998; Kawai and Akira 2010). These initial steps are followed by the activation of key transcription factors (e.g., NF-kB and interferon regulatory factors) that orchestrate the inflammatory and/or antiviral response signals involved in pathogen clearance and the subsequent development of appropriate adaptive immune responses (Medzhitov and Janeway 1998).

Although much attention has been devoted to characterizing transcriptional changes in response to infectious agents or other immune stimuli, we still know remarkably little about the contribution of post-transcriptional changes – specifically, changes in alternative pre-mRNA processing – to the regulation of the immune system (Lynch 2004; Martinez and Lynch 2013; Alasoo et al. 2015). Alternative splicing can impact cellular function by creating distinct mRNA transcripts from the same gene, which may encode unique proteins that have distinct or opposing functions. For instance, in human and mouse cells, several alternatively spliced forms of genes involved in the TLR pathway have been shown to function as negative regulators of TLR signaling in order to prevent uncontrolled inflammation (Rao et al. 2005; Wells et al. 2006; O’Connor et al. 2015). Additionally, several in-depth studies of individual genes suggest that alternative splicing plays an important role in increasing the diversity of transcripts encoding MHC molecules (Ishitani and Geraghty 1992) and in modulating intracellular signaling and intercellular interactions through the expression of various isoforms of key cytokines (Bihl et al. 2002; Nishimura et al. 2000), cytokine receptors (Rao et al. 2005; Goodwin et al. 1990; Koskinen et al. 2009), kinases (Jensen and Whitehead 2001) and adaptor proteins (Gray et al. 2010; Ohta et al. 2004; Janssens et al. 2002).

Yet, the extent to which changes in isoform usage are a hallmark of immune responses to infection remains largely unexplored at a genome-wide level (Alasoo et al. 2015; Rodrigues et al. 2013). Moreover, we still do not know which RNA processing mechanisms contribute most to the regulation of immune responses, or how these disparate mechanisms are coordinated.

To address these questions, we investigated genome-wide changes in transcriptome patterns after independent infections with *Listeria monocytogenes* or *Salmonella typhimurium* in primary human macrophages. Because of the distinct molecular composition of these two pathogens and the way they interact with host cells, they activate distinct innate immune pathways after infection (Haraga et al. 2008; Pamer 2004). Thus, our study design allowed us to evaluate the extent to which changes in isoform usage are pathogen-specific or more generally observed in response to bacterial infection. Furthermore, the large number of individuals in our study allowed us to use natural variation in RNA processing to gain insight into fine-tuned inter-individual regulation of immune responses. Our results provide a comprehensive picture of the role of RNA processing in regulating early innate immune responses to bacterial infection in human antigen-presenting cells.

## RESULTS

Infection with either *Listeria* or *Salmonella* induces dramatic changes in mRNA expression levels. Following 2 hours of infection with each bacteria, we collected RNA-seq data from 60 matched non-infected and infected samples, with an average of 30 million reads sequenced per sample (see Methods; Table S1). The first principal component of the complete gene expression data set clearly separated infected from non-infected samples, and both PC1 and PC2 clustered infected samples by pathogen (i.e. *Listeria* or *Salmonella;* Figure 1A). Accordingly, we observed a large number of differences in gene expression levels between infected and non-infected cells, with 5,809 (39%) and 7,618 (51%) of genes showing evidence for differential gene expression (DGE) after infection with *Listeria* and *Salmonella*, respectively (using DESeq2, FDR ≤ 0.1% and |log2(fold change) | > 0.5; Figure S1A, Table S2). As expected, the sets of genes that responded to either infection were strongly enriched (FDR ≤ 1.8×10^−6^) for genes involved in immune-related biological processes such as the regulation of cytokine production, inflammatory responses, or T-cell activation (Table S3).

**Figure 1.**
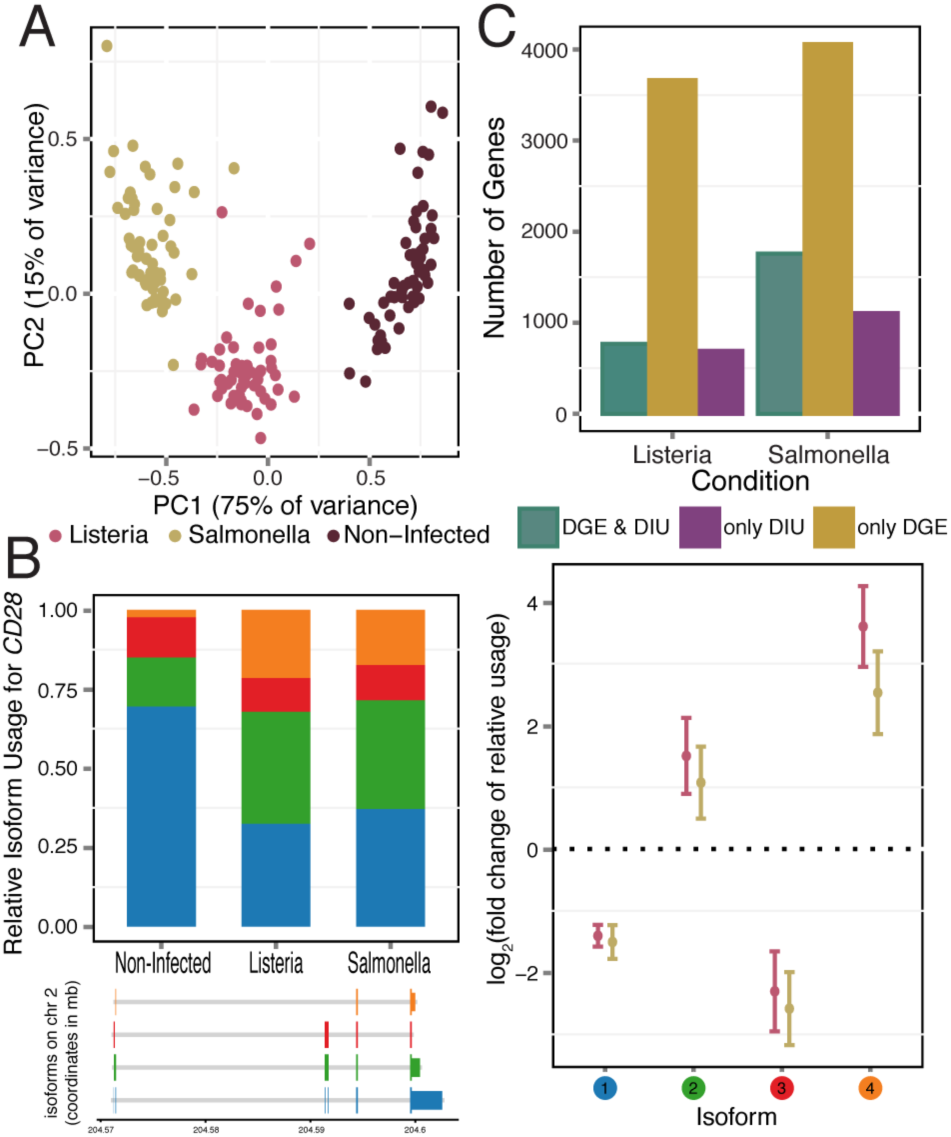
Gene expression and isoform proportion differences in response to bacterial infection. (A) Principal component analysis of gene expression data from all samples (PC1 and PC2 on the *x*- and *y-axis*, respectively). (B) *ADORA3*, a gene with significant changes in isoform usage upon infection with *Listeria* and *Salmonella*. For each *ADORA3* isoform in response to infection, plotted are the average relative isoform usages across samples (*left panel*) and corresponding fold changes (*right panel*; log2 scale; with standard error bars). Isoforms are ordered by relative abundance in non-infected samples, and colored circles (*right panel*) correspond to colors in barplot (*left panel*). (C) Number of genes showing only DIU, only DGE, and both DIU and DGE upon infection with *Listeria* and *Salmonella*, (11,353 genes tested).

In order to uncover changes in isoform usage in response to infection using our RNA-sequencing data, we applied two complementary approaches for analyzing RNA processing. We first measured abundances of full isoforms to gain a holistic perspective of the transcriptome landscape. Second, we used the relative abundances of individual exons within a gene to gain a finer understanding of the specific RNA processing mechanisms likely to be involved in the regulation of immune responses.

**Pervasive differential isoform usage in response to infection.** We initially sought to assess changes in isoform usage after infection using proportional abundances of the different isoforms encoded by the same gene. To do so, we used the transcripts per million (TPM) values reported by the RSEM software (Li et al. 2010) and calculated relative proportions by dividing isoform-level expression by the overall expression level of the gene (i.e., TPM summed across all possible isoforms). We then quantified differential isoform usage (DIU) between conditions for genes with at least two annotated isoforms (*N = 11,353*) using a multivariate generalization of the Welch's t-test that allowed us to test whether, as a group, the proportional abundances of the different isoforms in each gene were significantly different between infected and non-infected cells (Methods). We found that 1,456 (13%; after *Listeria* infection) and 2,862 (25%; after *Salmonella* infection) genes showed evidence for significant DIU after infection (FDR ≤ 1%; Figure 1B for an example of an DIU gene, Figure S1B, Table S4). Across all DIU genes, we observed a mean of 10% (± 8% s.d.) and 13% (± 10% s.d.) change in relative isoform usage upon infection with *Listeria* and *Salmonella*, respectively (defined as the maximum change upon infection in relative isoform usage across transcripts of each gene, Figure S1C). Following infection with *Listeria* and *Salmonella*, 6.2% and 10.2%, respectively, of genes with a dominant isoform prior to infection (see methods) switch to using an alternative dominant isoform (Figure S1E).

DIU genes were significantly enriched for genes involved in immune responses (FDR ≤ 0.01; Table S3), including several cytokines (e.g. *IL1B, IL7, IL20*), chemokines (*CCL15*), regulators of inflammatory signals (e.g. *ADORA3, MAP3K14*) and genes encoding co-stimulatory molecules required for T cell activation and survival (e.g. *CD28* in Figure 1B, *CD80*) (Figure S2). Notably, 86% of genes with DIU upon infection with *Listeria* were also classified as having DIU after infection with *Salmonella*, suggesting that changes in isoform usage are likely to be common across a wide variety of immune triggers. To assess the robustness of our findings, we tested for DIU using isoform-specific expression levels calculated with Kallisto, an alignment-free quantification method (Bray et al. 2016) (see Supplementary Methods). At an FDR of 5%, we observed a 74% (*Listeria*) and 78% (*Salmonella*) agreement with DIU genes identified using RSEM, confirming that identification of DIU genes is largely robust to the method used for isoform quantification.

Considering the large proportion of genes with significant DIU upon infection, we sought to understand whether these changes arose from a shift in the usage of a dominant isoform or increased isoform diversity within a given gene. To do so, we calculated the Shannon diversity index, which measures the evenness of the isoform usage distribution for each gene before and after infection (low values reflect usage of one or few isoforms; high values reflect either the usage of a more diverse set of isoforms or more equal representation of the same set of isoforms). Δ_shannon_ thus quantifies the change in isoform diversity after infection (Table S5). The majority of genes (63% in *Listeria* and 68% in *Salmonella*) show an increase in isoform diversity (Δ_shannon_ > 0) after infection (Figure S1D), indicating a shift away from usage of the primary (pre-infection) isoform after infection and a general increase in the diversity of transcript species following infection.

Previously, it had been reported that DGE and DIU generally act independently to shape the transcriptomes of different mammalian tissues (Barbosa-Morais et al. 2012). In contrast, we found that DIU genes were significantly enriched for genes with DGE (both up- and down-regulated genes, Fisher's exact test, *P* ≤ 7.5×10^−6^ for both *Listeria* and *Salmonella*) (Figure S3A). Despite such enrichment, a substantial proportion of genes (47% (690 genes) in *Listeria* infected samples and 39% (1,105 genes) in *Salmonella* infected samples) with significant changes in isoform usage after infection do not exhibit DGE following infection (Figure 1C). Even after relaxing the FDR threshold for DGE by 100-fold, 685 (*Listeria*) and 1096 (*Salmonella*) DIU genes still exhibited no significant DGE. Thus, a considerable fraction of transcriptome changes in response to infection do occur solely at the level of RNA processing, and independently of changes in mean expression levels.

**Directed shifts in 3' UTR length and alternative splicing following infection.** Though useful in documenting the substantial transcriptome changes upon infection, aggregate isoform levels calculated by RSEM do not distinguish between different types of RNA processing changes. Thus, we next examined changes in five distinct categories of alternative RNA processing, allowing us to zoom in on particular molecular mechanisms that underlie differential isoform usage in response to infection. Specifically, we used human transcript annotations from Ensembl to quantify usage of (1) alternative first exons (AFEs), (2) alternative last exons (ALEs), (3) alternative polyadenylation sites, leading to tandem 3’ untranslated regions (TandemUTRs), (4) retained introns (RIs), and (5) skipped exons (SEs). AFEs, ALEs, and TandemUTRs correspond to the alternative usage of terminal exons primarily affecting UTR composition, whereas RIs and SEs correspond to internal splicing events that usually affect the open reading frame.

For each gene or exon within each RNA processing category, we used the MISO software (Katz et al. 2010) to calculate a “percent spliced in” (PSI or Ψ) value (Table S6), defined as the proportion of transcripts from a gene that contain the “inclusion” isoform (defined as the longer isoform for RIs, SEs, and TandemUTRs, or use of the exon most distal to the gene for AFEs and ALEs). ΔΨ values thus represent the difference between PSI values calculated for the infected versus non-infected samples. Overall, we observed many significant changes in RNA processing (*N ≥ 1,098*) across all categories in both bacteria (significance was defined as Bayes Factor > 5 in at least 10% of individuals and |mean ΔΨ | > 0.05), compared to null expectations derived from measuring changes among pairs of non-infected samples (N=29) (Figure 2A, Table S7). Our criteria for determining significant changes also allows us to choose exons that are more likely to be consistently changing across many individuals; significantly changing exons have lower variance in ΔΨ values across individuals than exons that are not changing after infection (Figure S4). The greatest proportion of changes after infection occurred among retained intron and TandemUTR events. PSI values for 7.3% and 14% of retained introns and 7.6% and 16.7% of TandemUTRs significantly changed after infection with *Listeria* or *Salmonella*, respectively. When we considered the set of genes associated with at least one significant change in RNA processing, we observed an over-representation of Gene Ontology categories involved in immune cell processes and the cellular response to a stimulus (Figure 2B, Table S8).

**Figure 2.**
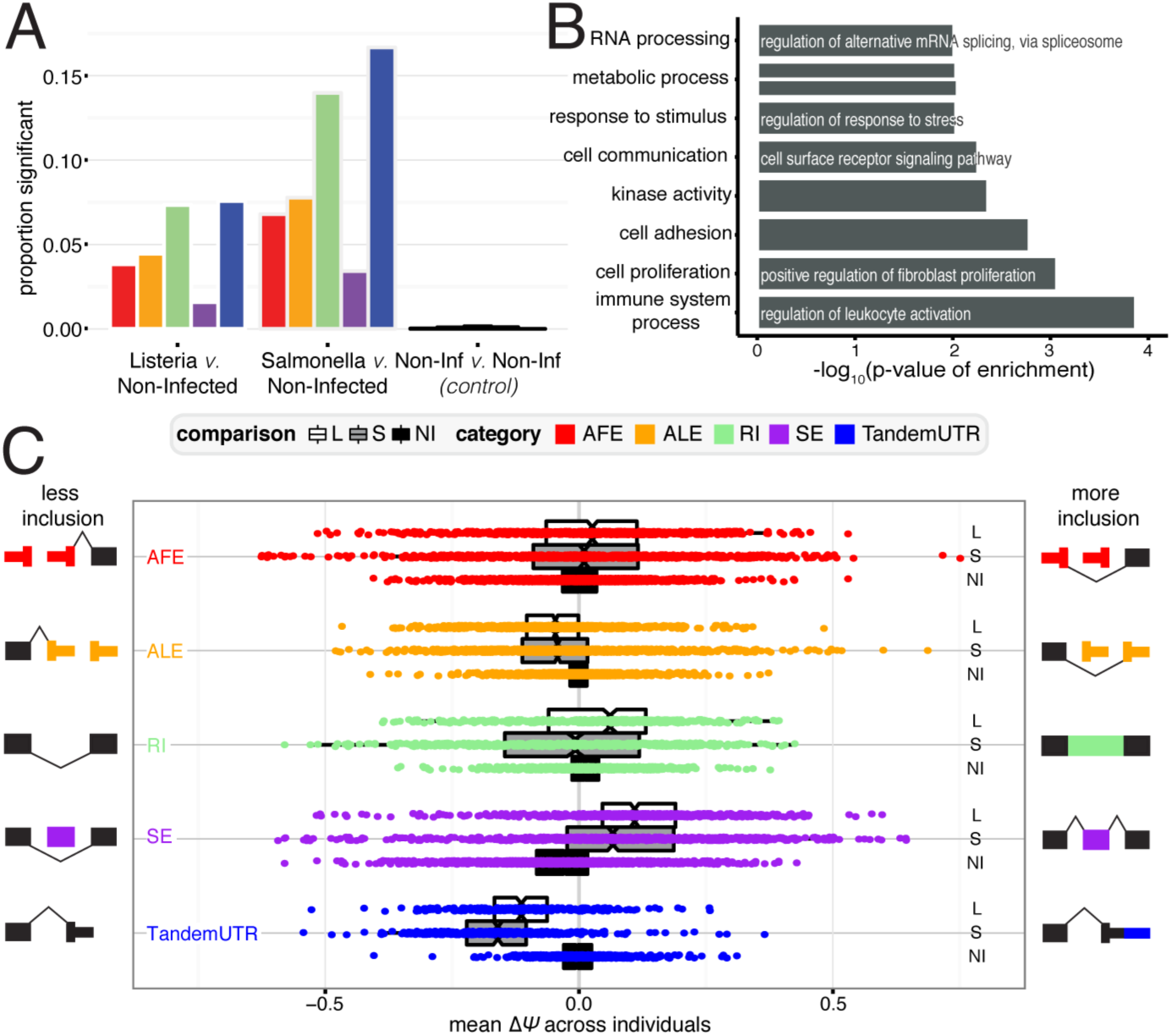
RNA processing changes in response to bacterial infection. Proportion of events for RNA processing category that are significantly changing after infection with *Listeria* (*left*), *Salmonella* (*middle*), or variation between noninfected samples as a control (*right*). Numbers indicate the number of significant changes per category. (B) Significantly Gene Ontology categories for genes with any significant RNA processing change (FDR ≤ 10%). (C) Distribution of ΔΨ values for each RNA processing category. Negative values represent less inclusion, while positive values represent more inclusion, as defined by the schematic exon representations.

Genome-wide shifts towards more prevalent usage of inclusion or exclusion isoforms have previously been observed as signatures of cellular stress responses or developmental processes (Sandberg et al. 2008; Mayr and Bartel 2009; Wong et al. 2013; Lackford et al. 2014a; Shalgi et al. 2014; Ji et al. 2009). To examine directional shifts in particular RNA processing mechanisms, we used mean ΔΨ values (defined as the mean ΔΨ across all individuals for each gene or exon) and observed a striking global shift towards the usage of upstream polyadenylation sites in tandem 3’ UTR regions, indicating pervasive 3’ UTR shortening following infection with either *Listeria* or *Salmonella* (Figure 2C; both *P* < 2.2×10^−16^ with Mann-Whitney U test; 94.4% and 98% of significant TandemUTRs changes in *Listeria* and *Salmonella*). Although less frequent, we also see a general trend towards usage of upstream polyadenylation sites in alternative last exons. Interestingly, we found a striking correlation in the extent to which a given individual has a global shift towards 3’ UTR shortening and towards usage of an upstream ALE (Pearson R = 0.83, *P* = 4.97 × 10^−16^ for Listeria and Pearson R = 0.78, *P* = 2.65 × 10^−13^ for Salmonella, Figure S6B). These correlations suggest that the changes in ALEs and TandemUTRs are likely being regulated by similar or coordinated mechanisms; rather they are likely regulated through changes in the activities of cleavage and polyadenylation factors (rather than splicing machinery), as suggested by previous reports (Takagaki and Manley 1998; Di Giammartino et al. 2011; Taliaferro et al. 2016). Additionally, we found a substantial increase in the inclusion of skipped exons after infection (both *P* < 2.2×10^−16^ with Mann-Whitney U test; 79.8% and 77.3 % of significant SE changes in *Listeria* and *Salmonella*, Figure S5). The consistency of these patterns suggests that a dedicated post-transcriptional regulatory program might underlie genome-wide shifts in RNA processing after bacterial infection.

Quantifying changes at the terminal ends of transcripts is challenging due to known biases in standard RNA-seq protocols (Wang et al. 2009). Thus, we validated our observation of 3’ UTR shortening upon infection by sequencing RNA from 6 individuals after infection with *Listeria* and *Salmonella* with a 3’RNA-seq protocol, which specifically captures the 3’ most end of transcripts (see Methods). TPM expression values calculated from 3’ RNA-seq data highly correlate with TPMs from RNA-seq data, establishing that this method is able to quantitatively measure of the number of transcripts in a cell (R > 0.82 across all conditions, *P* < 2.2 × 10^−16^, Figure S6). Meta-gene plots around the polyadenylation sites of core and extended regions of TandemUTRs show a notable increase after infection in sequencing coverage around the upstream polyadenylation site relative to the downstream polyadenylation site, specifically for genes with significantly shorter 3’ UTRs upon infection as assessed by the RNA-seq data (Figure 3A) Finally, to look at 3’ UTR shortening in specific genes, we calculate Ψ values for TandemUTRs using the 3’RNA-seq data (see Supplementary Methods) and find again that genes that are significantly changing 3’ UTR usage in the RNA-seq data exhibit a significant shift towards negative ΔΨ values in the 3’RNA-seq data (P < 2.2 × 10^−16^; Figure 3B).

**Figure 3.**
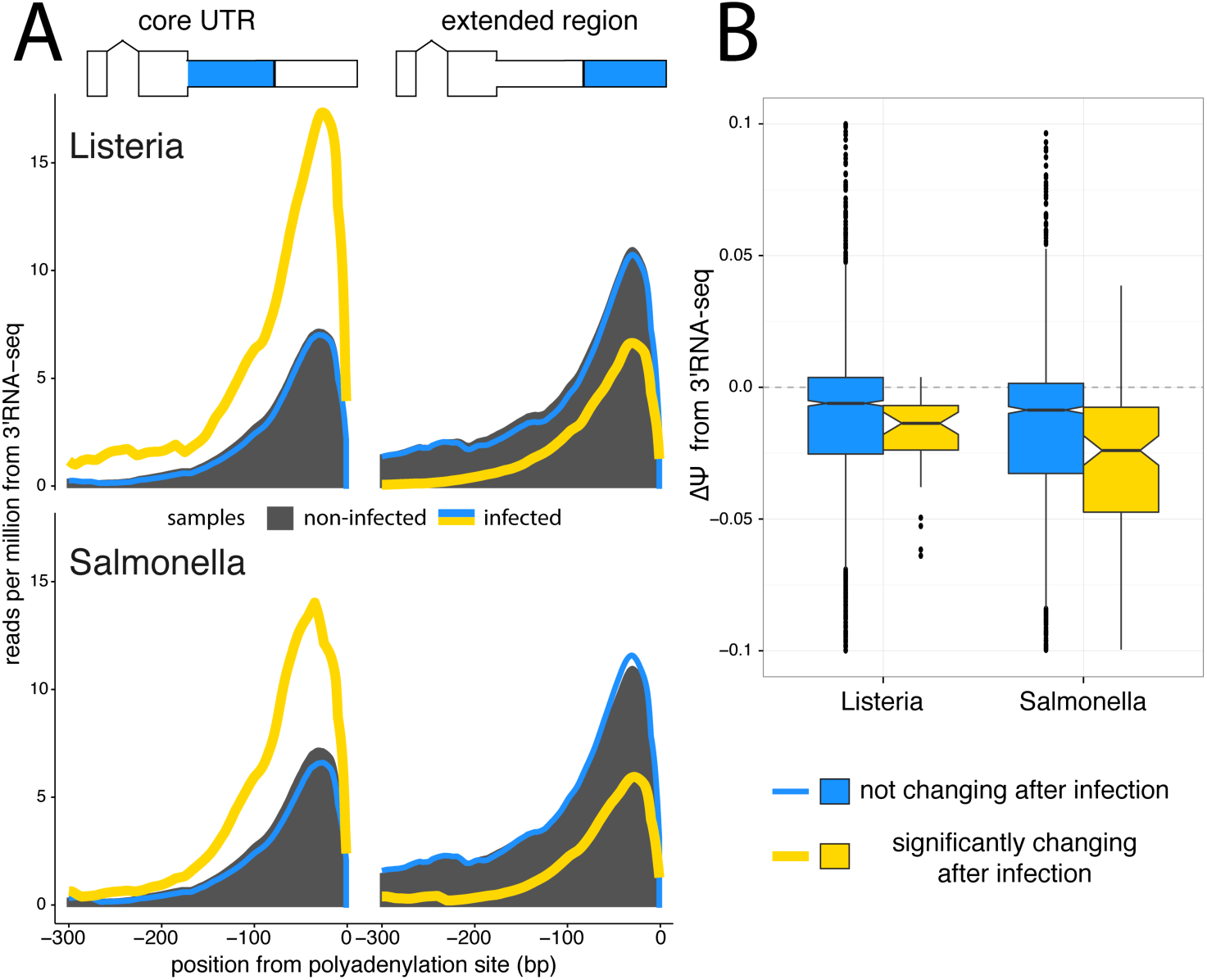
3’RNA sequencing shows increased usage of upstream polyadenylation sites upon infection. (A) Meta-gene distributions of 3’RNA-seq read densities at the upstream polyA sites (core regions, *left*) and downstream polyA sites (extended regions, *right*) of Tandem 3’ UTRs after infection with *Listeria* or *Salmonella (top* and *bottom*, respectively). Shown are the read distributions for non-infected samples across all Tandem 3’ UTRs (*black*) and infected samples at Tandem 3’ UTRs that significantly change after infection (*yellow*) or show no change after infection (*blue*), as called by the RNA-seq data. (B) Distribution of ΔΨ values calculated from 3’RNA-seq data for Tandem 3’ UTRs. We observe significant shifts (*P* < 2.2 × 10^−16^ for both *Listeria* and *Salmonella*) towards negative ΔΨ values in Tandem UTRs that are identified as significantly changing in RNA-seq data (*yellow*) relative to Tandem UTRs without any change after infection (*blue*).

**RNA processing changes persist 24 hours after infection.** To evaluate whether the strong global shifts we observed were the result of a sustained immune response to the bacteria rather than a short-lived cellular stress response, we sequenced RNA from a subset of our individuals (*N*=6) after 24 hours (h) of infection with *Listeria* or *Salmonella*. Samples sequenced after 24h were paired with matched non-infected controls cultured for 24h to calculate ΔΨ values for alternative RNA processing categories using MISO (Katz et al. 2010). Across all categories, we found many more significant changes in RNA processing and larger ΔΨ values after infection for 24h than after 2h (Figure S7). These results further support a key role for RNA processing in controlling innate immune responses to infection. For genes or exons that changed significantly after 2h of infection in this set of 6 individuals (see Methods), the majority of changes (67%) remained detectable at 24h, usually with similar magnitude and direction (Figure 4A). The one notable exception is that introns that are preferentially retained at 2h in infected cells tended to be spliced out at 24h. Our results thus indicate that the splicing of alternative retained introns during the course of the immune response to infection is temporally dynamic.

**Figure 4.**
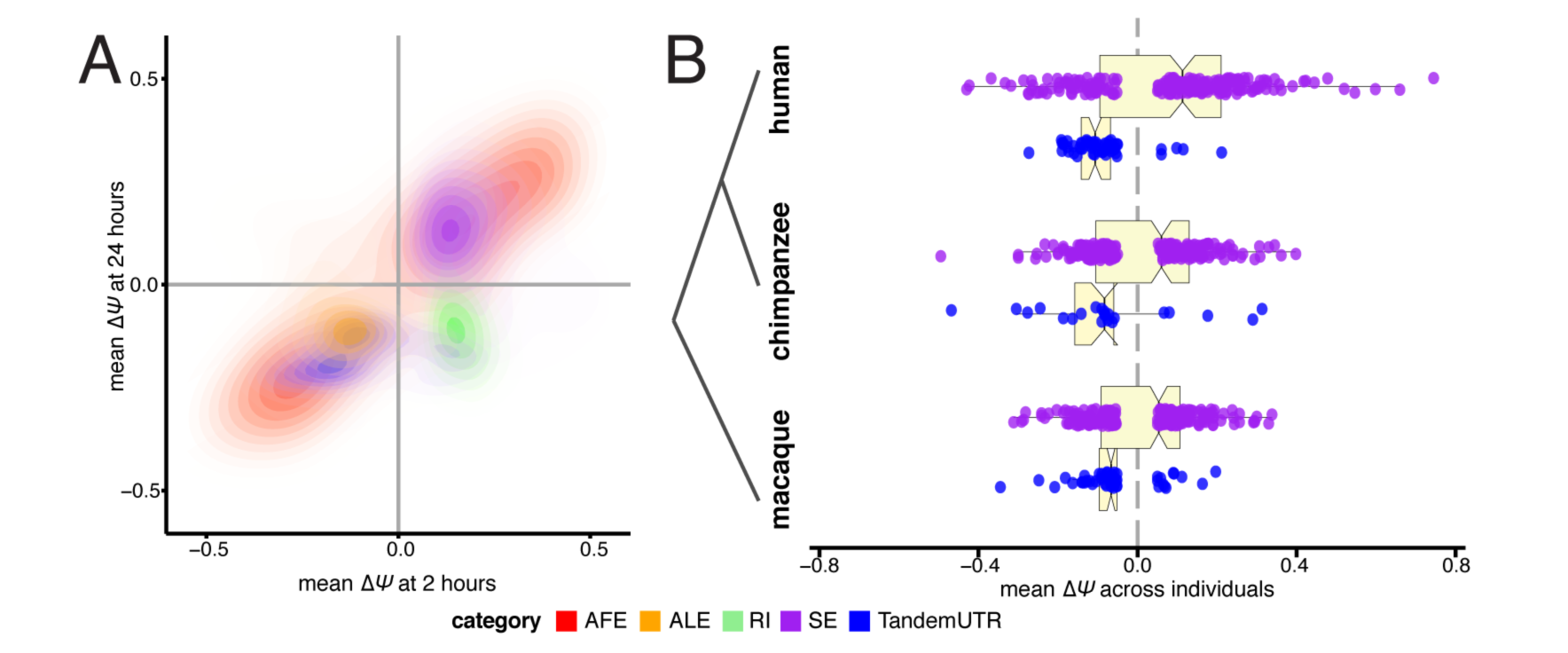
Directed shifts in RNA processing persist across time, stimulus conditions, and closely related species. (A) Correlations between ΔΨ values after 2 hours of infection (*x-axis*) and ΔΨ values after 24 hours of infection (*y-axis*), presented as density plots per RNA processing category in contrasting colors. Plotted are events that are significant after 2 hours of either *Listeria* or *Salmonella* infection. (B) Distributions of ΔΨ values for skipped exons (*purple*) and TandemUTRs (*blue*) following infection of whole blood cells with lipopolysaccharide, assessed in human (*top*), chimpanzee (*middle*) and rhesus macaque (*bottom*) individuals (*N=6* per species). We observe prominent global shifts in isoform distributions in human and macaque (SE_human_ *P=0.003*, TandemUTR_human_ *P=2.4×10^−13^;*, SE_macaque_ *P=0.05*, TandemUTR_macaque_ *P=8.8×10^−6^*) with a more modest trend observed in chimpanzee, potentially due to poor transcript annotations in the chimpanzee genome (SE_chimpanzee_ *P=0.26*, TandemUTR_chimpanzee_ *P=0.002*). All p-values were calculated using a Student’s t-test testing deviation from a mean of zero.

**Shortening of 3' UTR regions and increased exon inclusion are evolutionarily conserved responses to infection in primates.** Since the innate immune response is ancient, one might expect core aspects of this pathway to be evolutionarily conserved among primates. To explore this question, we used high-depth RNA-seq data from human, chimpanzee, and rhesus macaque primary whole blood cells (*N*=6 individuals from each species) before and after 4 hours of stimulation with lipopolysaccharide (LPS), the major component of the outer membrane of Gram-negative bacteria. Using an analysis pipeline similar to our analysis of infection after 24 hours of infection with *Listeria* or *Salmonella*, we calculated ΔΨ values for alternative RNA processing categories using MISO (Katz et al. 2010) (see Supplementary Methods, Table S9). As with macrophages, we observed consistent strong shifts towards increased exon inclusion and shorter 3’ UTRs in human whole blood cells stimulated with LPS. Furthermore, we observed similar trends toward increased exon inclusion and significant 3' UTR shortening in macaques and chimpanzees, supporting a conserved response of RNA processing pathways during innate immune responses in primates (Figure 4B). In addition, these data shows that the shortening of 3’ UTRs and increased exon inclusion are not limited to isolated macrophages but also observed as part of the immune response engaged by interacting primary granulocytes and lymphocytes, more closely mimicking *in vivo* cellular responses to infection.

**Increased exon inclusion is associated with increased gene expression levels.** Since we observed strong connections between overall isoform usage and differential gene expression, we explored the relationship between the occurrence of particular RNA processing changes and the direction of gene expression changes in response to infection. We found that genes with significant differences in skipped exon usage pre- and post-infection (up to 80% of which have increased skipped exon inclusion after infection) were more likely to be up-regulated after infection (T-test, both *P* ≤ 1×10^−4^ for *Listeria* and *Salmonella;* Figure 5A; Figure S8A). This association remained when gene expression levels were calculated using reads mapping to constitutive exons only (Figure S8A), eliminating concerns about alternative spliced isoforms impacting gene expression estimates. Our observation is consistent with a recent study that reported increased gene expression in tissues with increased inclusion of evolutionarily novel SEs (Merkin et al. 2015). Genes with other types of splicing changes showed no trend in expression changes.

**Figure 5.**
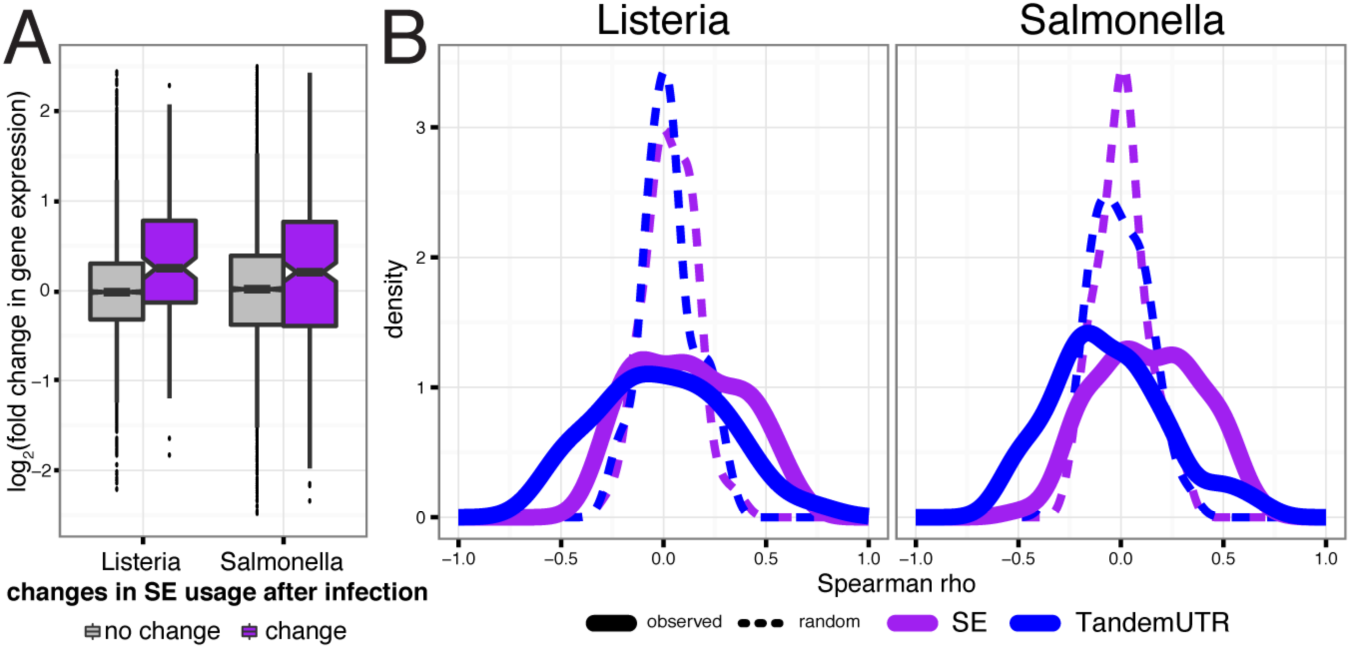
Relationship between RNA processing changes and gene expression changes. (A) Distribution of fold changes in gene expression (*y-axis*, log2 scale) for genes with significant skipped exon changes after infection (*purple*) and genes with no change after infection (*gray*). (B) Distribution of Spearman correlations between ΔΨ and fold change in gene expression across 60 individuals per gene (*solid line*), for genes with only one annotated alternative event and significantly changed SE usage (*purple;* n_L_ = 46 genes and n_S_ = 97 genes) or tandem UTR usage (*blue;* n_L_ = 36 genes and n_S_ = 86 genes). Dotted lines show distribution of the correlation coefficients after permuting ΔΨ values.

Genes with significant skipped exon usage were enriched for Gene Ontology categories such as “response to stimulus,” and also for “RNA processing” and “RNA stability” categories (Figure S9A). Included within these latter categories are several members of two major splicing factor families, SR proteins and hnRNPs, which showed increased inclusion of skipped exons and increased gene expression after infection (Figure S9B). The SR protein and hnRNP families are known to function in complex auto- and cross-regulatory networks (Huelga et al. 2012; Pandit et al. 2013), often antagonizing one another’s effects on splicing (House and Lynch 2008; Wang and Burge 2008; Busch and Hertel 2012). Consistent with the slightly higher up-regulation of members of the hnRNP family, we observed a consistent enrichment for intronic splicing silencers in upstream intronic regions around significantly excluded skipped exons (Figure S9C). These elements are often bound by hnRNPs to promote exclusion of a cassette exon (Wang and Burge 2008))

**Variation across individuals provides insight into putative trans-regulatory factors.** A unique feature of our study design is our sampling of 60 individuals after infection with both *Listeria* and *Salmonella*. By taking advantage of natural variation in the regulation of RNA processing to infection, we aimed to gain further insight into the connections between RNA processing and changes in gene expression levels in response to infection. For each gene, we calculated the correlation between the inter-individual variation in RNA processing changes (ΔΨ) and the fold changes in gene expression levels upon infection. For most categories, we observed shifts in the distribution of correlations between RNA processing and gene expression levels relative to permuted controls (Figure S8B), with skipped exons and TandemUTRs showing the most consistent patterns (Kolmogorov-Smirnov test, *P* ≤ 0.005; Figure 5B). Increased skipped exon inclusion correlates with increased gene expression of the gene across individuals in 69% of the genes with significant skipped exons. In addition, preferential expression of shorter alternative 3’ UTR isoforms tends to be correlated with increased up-regulation of the associated genes in response to infection. Our findings thus suggest that RNA processing changes may directly impact gene expression levels, or at least are commonly regulated with changes in transcription or mRNA stability.

Substantial inter-individual variation in the genome-wide average RNA processing patterns also correlated strongly with average gene expression changes upon infection, particularly for changes in 3’ UTR usage. That is, individuals who exhibited more skipped exon inclusion or 3’ UTR shortening also tended to exhibit larger global shifts towards up-regulation of gene expression levels upon infection (Figure S10B).

Next, we sought to exploit our relatively large sample size to identify candidate *trans*-factors impacting the amount of skipped exon and TandemUTR changes observed across individuals. Specifically, we examined the relationship between mean changes in exon inclusion and fold change in expression for all expressed genes, across the 60 individuals. When we focus on proteins with known RNA binding functions or domains – hypothesizing that these factors are more likely to regulate post-transcriptional mechanisms – we find many significant correlations (FDR ≤ 1%) for known regulators of the corresponding RNA processing category (Figure S11). For instance, genes whose changes in expression correlated with individual-specific mean 3’ UTR shortening included several factors previously implicated in regulating polyadenylation site usage (such as cleavage and polyadenylation factors (Zheng and Tian 2014), hnRNP H proteins (Katz et al. 2010), and hnRNP F (Veraldi et al. 2001); Figure S11A). Additionally, many known regulators of alternative splicing – including a number of SR proteins and hnRNPs – have increased expression levels in individuals with larger extents of skipped exon inclusion (Figure S11B).

**3' UTR shortening as a means to evade repression by immune-related miRNAs.** The largest global shift observed in alternative isoform abundance was for 3’ UTR shortening, where as many as 98% of significantly changing TandemUTRs shifted towards usage of an upstream polyA site post-infection. Previously, global 3’ UTR shortening has been associated with proliferating cells, particularly in the context of cellular transitions in development (Sandberg et al. 2008), cell differentiation (Lackford et al. 2014b), cancerous (Mayr and Bartel 2009) states, or global 3’ UTR lengthening in embryonic development (Ji et al. 2009). However, macrophages do not proliferate, which we confirmed with a BrdU labeling assay in both resting and infected conditions (Figure S12). This result indicates that the pervasive 3’ UTR shortening observed in response to infection is due to an active cellular process that is independent of cell division and might be mechanistically distinct from that leading to 3’ UTR shortening in proliferating cells.

Previous studies in proliferating cells postulated that 3’ UTR shortening can act as a way to evade regulation by microRNAs (miRNAs), since crucial miRNA target sites are most often found in 3’ UTR regions (Sandberg et al. 2008). To evaluate this hypothesis we performed small-RNA sequencing in 6 individuals after 2h and 24h of infection with *Listeria* and *Salmonella*. When focusing specifically on miRNAs expressed in macrophages (Table S10), we found that the extended UTR regions of infection-sensitive 3’ UTRs have a significantly higher density of miRNA target sites compared to TandemUTRs that do not change in response to infection (Figure 6A). This increased density of miRNA target sites was restricted to the extension region, which is subject to shortening after infection: no difference in the density of target sites was observed in the common “core” 3’ UTR regions of the same genes (Figure S13).

**Figure 6.**
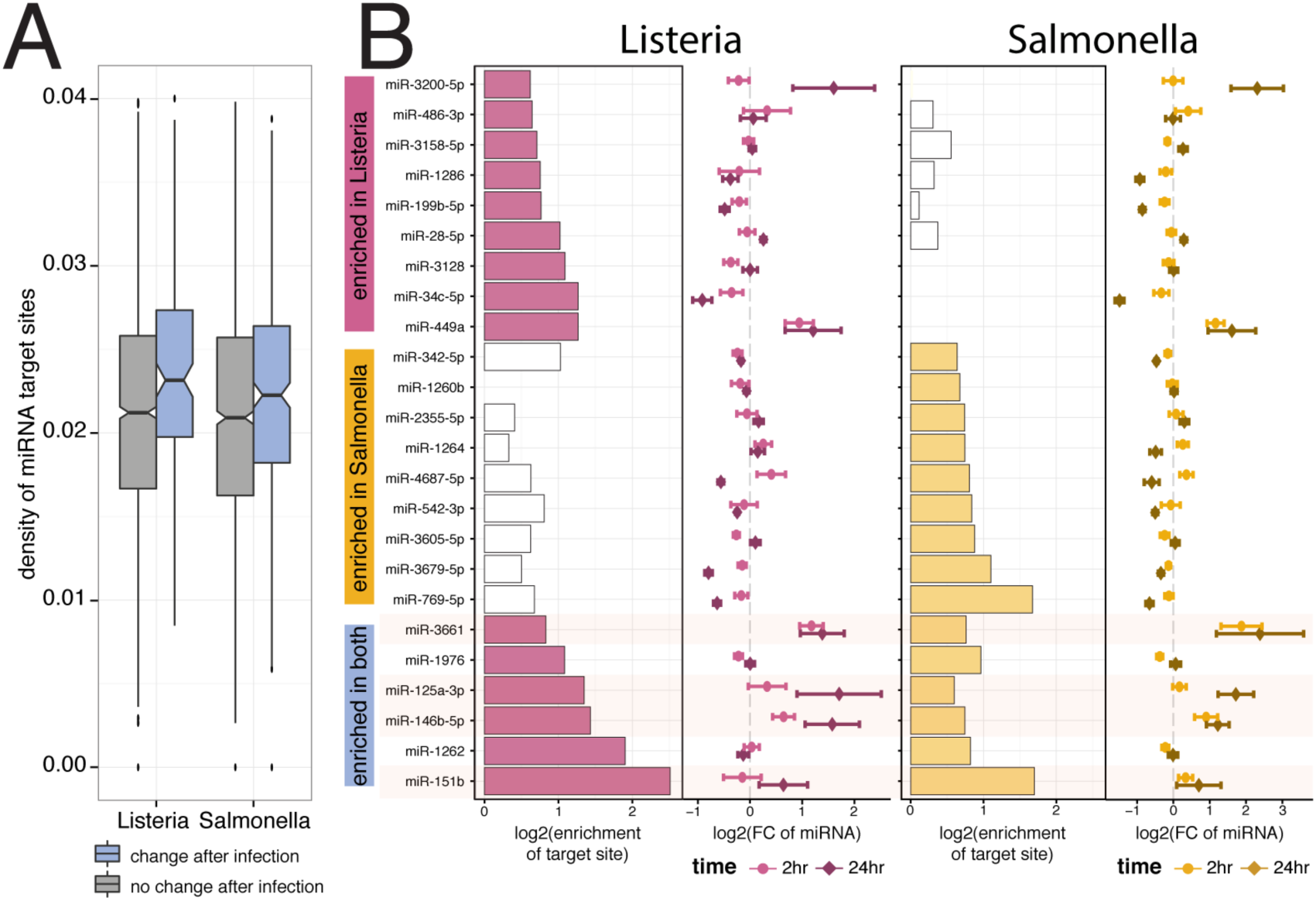
Tandem 3’ UTR shortening allows evasion of regulation by miRNAs. (A) Distribution of frequency of miRNA target sites per nucleotide in the extended regions of Tandem UTRs that either show no change after infection (*grey*) or significantly change after infection (*blue*). (B) Significantly enriched miRNA target sites (FDR ≤ 10%, |FC | > 1.5) in the extended regions of significantly changing Tandem UTRs after infection with Listeria-only (top), with Salmonella-only (middle), or both bacteria (bottom). For each bacteria, the barplots in the left panels show the fold enrichment (*x-axis*, log_2_ scale) of target sites in the extended regions. White bars represent non-significant enrichments. Panels on the right show the fold change in miRNA expression (*x-axis*, log_2_ scale with standard error bars) after either 2 hours of infection (*light colors*) or 24 hours of infection (*dark colors*).

Next, we tested if the increased density of miRNA target sites was driven by the enrichment of target sites for particular miRNAs expressed in macrophages. For each miRNA that was expressed in either non-infected or infected macrophages, we calculated an enrichment score assessing whether the target sites of that miRNA were significantly enriched in the extended region of significantly shortened 3’ UTRs (with a background distribution matched for sequence composition; Figure S14; see Supplemental Methods). We found 15 miRNAs with significantly enriched target sites (FDR ≤ 10% and enrichment ≥ 1.5 fold) in either the *Listeria* or *Salmonella* conditions, with 6 miRNAs over-represented in both bacterial conditions (Figure 6B). Interestingly, of the miRNAs with the highest enrichment after infection with either *Listeria* or *Salmonella*, two (miR-146b and miR-125a) have previously been shown to be important regulators of the innate immune response (Taganov et al. 2006; Perry et al. 2009; Boldin and Baltimore 2012; Kim et al. 2012; Banerjee et al. 2013). Accordingly, we found that 4 of the miRNAs with the highest enrichment after infection (miR-125a, miR146b, miR-3661, and miR-151b) are all up-regulated following infection, and strongly so after 24h of infection (Figure 6B). In the majority of cases, greater 3’ UTR shortening was associated with a stronger increase in gene expression across individuals, suggesting that shifts towards shorter 3’ UTRs after infection allows these transcripts to escape from repression by specific immune-induced miRNAs.

## DISCUSSION

Taken together, our results provide strong evidence that RNA processing plays a key role in the regulation of innate immune responses to infection. Despite known differences in the regulatory pathways elicited in macrophages by *Listeria* and *Salmonella*, our data revealed striking similarities in the overall patterns of RNA processing induced in response to both pathogens. Given the strong overlaps, we chose throughout this study to focus on consistent patterns across the two bacteria.

Though genes that have differential isoform usage in response to infection are more likely to show changes in gene expression, we found that as many as 47% of genes showing differential isoform usage have no evidence for differences in gene expression upon infection. This observation suggests that a considerable proportion of genes are regulated predominantly by post-transcriptional mechanisms, with minimal changes in transcriptional regulation. Differential isoform usage is coupled to a prominent increase in the diversity of isoforms following infection. Intriguingly, genes showing the strongest increases in diversity upon infection were strongly enriched among down-regulated genes (Figure S3B). This observation raises the possibility that the increased isoform diversity could reflect a shift toward less stable isoforms, e.g., those targeted by nonsense-mediated decay (NMD), thus reducing transcript levels (Lewis et al. 2003; Green et al. 2003).

Upon infection of human macrophages with either *Listeria* or *Salmonella*, we observed substantial shifts towards increased skipped exon inclusion and shortening of 3’ UTRs. These RNA processing patterns remained prominent when interrogated in human whole blood after an alternative immune stimulus (LPS), despite the fact the number of significant RNA processing changes in whole blood was generally smaller than that observed in purified macrophages infected with live bacteria. This reduced effect is likely explained by the fact that in whole blood, we are interrogating a very heterogeneous cell population within which only a fraction (~3–6%) of the cell types are actually responding to LPS stimulation, such as monocytes/macrophages (Zarember and Godowski 2002). Additionally, transcriptional changes induced by live bacteria such as *Listeria* or *Salmonella*, which concomitantly activates multiple innate immune pathways (Haraga et al. 2008; Pamer 2004), are grander than transcriptional changes that can induced by a single purified ligand, such as LPS, which activates primarily the TLR4 pathway (Poltorak et al. 1998). Despite this reduced statistical power, we still observed conserved RNA processing signatures of immune stimulation through the primate lineage, supporting the likely functional importance of these cellular responses and the ubiquity of RNA processing changes after immune stimulation across evolution.

The most prominent hallmark of RNA processing changes after infection is a global shortening of 3’ UTRs. Interestingly, this effect resembles the global 3’ UTR shortening seen in proliferating activated T cells and transformed cells (Sandberg et al. 2008) (Mayr and Bartel 2009) (Lackford et al. 2014b). Given that both T-cell and macrophage activation result in the activation of shared signaling cascades (e.g., c-Jun N-terminal kinase (JNK) – a key regulator of apoptosis – and nuclear factor κ-B (NF-κB) activation), it is possible that, at least to some extent, 3’ UTR shortening observed in T-cells and macrophages stems from the activation of these shared pathways. Previous studies hypothesized that 3’ UTR shortening allows mRNAs to escape regulation by dynamically changing miRNAs, suggesting that macrophages may co-opt generally used mechanisms for the same purpose. (Sandberg et al. 2008; Mayr and Bartel 2009). This shift towards shorter 3’ UTRs could result from increased miRNA-dependent degradation of transcripts with longer 3’ UTRs postinfection. However, genes that shift towards shorter 3’ UTRs were more highly expressed post-infection, making this explanation unlikely. Furthermore, the increase in overall miRNA target site density is reminiscent of a previous observation that genes with regulated 3’ UTR usage across tissues have more target sites for ubiquitously expressed miRNAs in their distal 3’ UTRs (Lianoglou et al. 2013). However, in contrast to results across tissues, we observe that the target sites in our distal 3’ UTR regions are directly related to miRNAs whose expression specifically changes after infection.

Specifically, our data support previous hypotheses and find evidence for that 3’ UTR shortening in macrophages is associated with the loss of target sites for a subset of immune-regulated miRNAs, including miR-146b, miR-125a, and miR-151b (Figure 6B). These miRNAs target the extended 3’ UTR regions of several critical activators of the innate immune response – including the *IRF5* transcription factor (Lawrence and Natoli 2011) and the MAP kinases *MAPKAP1* and *MAP2K4* (Arthur and Ley 2013) – all of which shift to usage of shorter 3’ UTRs after infection. It is tempting to speculate that without such 3’ UTR shortening, the binding of one or more of miRNAs to the 3’ UTR region of these genes would compromise their up-regulation and the subsequent activation of downstream immune defense pathways. Given that many miRNAs are involved in either (or both) mRNA degradation and translational efficiency, we cannot know for sure which of these processes is prevented by the 3’ UTR shortening of these genes.

The convergence towards similar isoform outcomes across many disparate genes suggests the activation of factors acting in *trans* to drive global shifts towards inclusion of cassette exons or usage of upstream proximal polyadenyalation sites. Taking advantage of our relatively large sample size, we were able to identify candidate *trans*-factors whose up-regulation upon infection might have widespread impacts on RNA processing patterns, including some known regulators of alternative splicing or 3’end processing (Figure S11). Sets of RNA binding factors likely act in combination to influence final transcriptome states (Brooks et al. 2015). For instance, a general up-regulation of splicing machinery following infection (Munding et al. 2013), allowing the cell to recognize and use additional splice sites, might contribute to increased exon inclusion and greater isoform diversity observed following infection. Members of both the hnRNP and SR protein splicing factor families show up-regulation of their overall gene expression levels (Figure S9B), likely contributing to specific instances of exon inclusion as well as exclusion. Correspondingly, changes in the regulation of many canonical cleavage and polyadenylation factors were tied to individual-specific shifts in 3’ UTR shortening. The significant correlation between global shifts in ALE usage and TandemUTRs across individuals further support the notion that cleavage and poladenylation pathways are regulating the changing 3’ UTR landscape upon infection (Figure S10A). Intriguingly, fold changes in hnRNP H1 gene expression have high correlations with both skipped exon inclusion and 3’ UTR shortening, consistent with its known splicing function and suggesting that this factor may play multiple roles in shaping the transcriptome in response to bacterial infection.

An alternative mechanism that has been proposed for the global regulation of 3’ UTR shortening in T-cell activation and neuronal differentiation is a phenomenon called *telescripting*. This is a phenomenon by which moderately lower U1 snRNA levels relative to global increases in nascent mRNA production restrict U1’s secondary role in inhibiting premature cleavage and polyadenylation (Berg et al. 2012). While we did not directly measure mRNA levels, we observe a moderate but significant increase in total RNA concentration after 2 hours of infection that is consistent with the increase in nascent RNA observed by Berg et al. (Berg et al. 2012) (across 60 samples, paired t-test; P=0.006 for Listeria and P=4.57 × 10^−6^ for Salmonella), combined with no changes in the levels of U1 snRNA (relative to 5S rRNA, Supplemental Methods, Figure S15). Thus it is possible that a delayed response of U1 snRNA could promote the usage of upstream polyadenylation sites in both TandemUTRs and ALEs. Interestingly, Berg et al. also observe increased up-regulation of individual internal exons upon moderate U1 knockdown in neurons (Berg et al. 2012), which is consistent with our observation of increased skipped exon inclusion.

Substantial inter-individual variation in the genome-wide average RNA processing patterns also correlated strongly with average gene expression changes upon infection, particularly for changes in 3’ UTRs and skipped exons. That is, individuals who exhibited more skipped exon inclusion or usage of upstream polyadenylation sites also tended to exhibit larger global shifts towards up-regulation of gene expression levels upon infection (Figure S10B). Though these patterns might result from stochastic variation in levels of one or several trans-factors, they are also likely to be influenced by genetic differences among individuals. It will be interesting to ask whether global isoform differences might potentially be reflective of variation in an individual’s susceptibility to infection.

### Materials and Methods

Complete details of the experimental and statistical procedures can be found in SI Materials and Methods. Briefly, blood samples from 60 healthy donors were obtained from Indiana Blood Center with informed consent and ethics approval from the Research Ethics Board at the CHU Sainte-Justine (protocol #4022). All individuals recruited in this study were healthy males between the ages of 18 and 55 y old. Blood mononuclear cells from each donor were isolated by Ficoll-Paque centrifugation and blood monocytes were purified from peripheral blood mononuclear cells (PBMCs) by positive selection with magnetic CD14 MicroBeads (Miltenyi Biotec). In order to derive macrophages, monocytes were then cultured for 7 days in RPMI-1640 (Fisher) supplemented with 10% heat-inactivated FBS (FBS premium, US origin, Wisent), L-glutamine (Fisher) and M-CSF (20ng/mL; R&D systems). After validating the differentiation/activation status of the monocyte-derived macrophages we infected them at a multiplicity of infection (MOI) of 10:1 for *Salmonella typhimurium* and an MOI of 5:1 for *Listeria monocytogenes* for 2- and 24-hours. For primate comparisons, whole blood was drawn from 6 healthy adult humans (CHU Sainte-Justine), 6 common chimpanzees, (Texas Biomedical Research Institute), and 6 rhesus macaques (Yerkes Regional Primate Center). Human samples were acquired with informed consent and ethics approval from the Research Ethics Board (CHU Sainte-Justine, #3557). Non-human primate samples were acquired in accordance with individual institutional IACUCC requirements. For all species, 1ml of blood was drawn into a media-containing tube (Trueculture tube, Myriad, US) spiked with ultrapure LPS (Invitrogen, USA) or endotoxin-free water (control). Samples were stimulated with 1ug/ml LPS, at 37C for 4 hours before total blood leukocytes were isolated and RNA collected. Genome-wide gene expression profiles of untreated and infected/treated samples were obtained by RNA-sequencing for both mRNA transcripts and small RNAs. After a series of quality checks (SI Materials and Methods), mRNA transcript abundances were estimated using RSEM (Li and Dewey 2011) and Kallisto (Bray et al. 2016) and microRNA expression estimates were obtained as previously described(Siddle et al. 2014). To detect genes with differential isoform usage between two groups of non-infected and infected samples we used a multivariate generalization of the Welch's t-test. Shannon entropy H_sh_ (also known as Shannon index) was applied to measure the diversity of isoforms for each target gene before and after infection. Changes across individual RNA processing events were quantified using the MISO software package (v0.4.9) (Katz et al. 2010) using default settings and hg19 version 1 annotations. Events were considered to be significantly altered post-infection if at least 10% (n >= 6) of individuals had a BF >= 5 and the |mean ΔΨ | ≥ 0.05 (Table S6). To conduct a targeting sequencing of 3’ ends of transcripts, We used Smart-3SEQ (Foley et al, manuscript in preparation), a modified version of the 3SEQ method (Beck et al. 2010). We used a custom gene ontology script to test for enrichment of functional annotations among genes that significantly changed isoform usage in response to infection (SI Materials and Methods). All predicted miRNA target sites within annotated TandemUTR regions were obtained using TargetScan (v6.2) (Friedman et al. 2009).

### Data Access

Data generated in this study have been submitted to the NCBI Gene Expression Omnibus (GEO; http://www.ncbi.hlm.nih.gov/geo/) under the accession number GSE73765, which comprises of the mRNA-seq data (GSE73502) and small RNA-seq data (GSE73478).

## Acknowledgements

We thank P Freese for a categorized list of RNA binding proteins, G McVicker for an initial script to perform iterative gene ontology analyses and P Sudmant for help parsing primate transcript annotations. We thank Y Katz, JM Taliaferro, P Sudmant, J Tung, and members of the Barreiro and Burge labs for helpful discussions and comments on the manuscript. This work was supported by grants from the elife senio (232519), the Human Frontiers Science Program (CDA-00025/2012) and the Canada Research Chairs Program (950-228993) (to L.B.B). We thank Calcul Québec and Compute Canada for managing and providing access to the supercomputer Briaree from the University of Montreal, used to do many computations reported in the manuscript. Additional computational resources were performed on an MIT computational cluster supported by the National Science Foundation (Grant No. 0821391). A.A.P. was supported by a Jane Coffin Childs postdoctoral fellowship. G. B. was supported by a postdoctoral fellowship from the Fonds de la Recherche en Santé du Québec.

